# Competitive binding of independent extension and retraction motors explains the quantitative dynamics of type IV pili

**DOI:** 10.1101/2020.02.12.946426

**Authors:** Matthias D. Koch, Chenyi Fei, Ned S. Wingreen, Joshua W. Shaevitz, Zemer Gitai

## Abstract

The functions of type IV pili (TFP) are mediated by cycles of extension and retraction. The coordination of these cycles remains mysterious due to poor quantification of TFP dynamics. Here we fluorescently label the TFP in the opportunistic pathogen *Pseudomonas aeruginosa* and track the full extension and retraction cycles of individual TFP to quantify their dynamics. We test several models for the switch between extension and retraction using quantitative experiments, biophysical modeling and genetics. We invalidate the prominent hypothesis that this switch is triggered by surface contact. Instead, we show that the entire repetitive cycle of extension and retraction of individual TFP is governed by the stochastic binding of antagonistic extension and retraction motors and explain how this mechanism quantitatively defines physiologically-important features like TFP length and their production rate. Interestingly, our results suggest that the major throttle of TFP production is the unbinding of the retraction motor.

## Introduction

Type IV pili (TFP) are amazing molecular machines that extend and retract extracellular polymers used for many biological functions^1–3^. TFP have emerged to be of particular interest in the opportunistic human pathogen *Pseudomonas aeruginosa*, as they promote surface motility, colonization, biofilm formation, and surface sensing^4–11^. In *P. aeruginosa*, the semi-flexible polymers of TFP are based on the major pilin (PilA) subunits whose extension is mediated by the PilB molecular motor and whose retraction is mediated by the PilT motor^2,3^. The structures of TFP and the components that build them have been well characterized by static methods such as electron microscopy^12^. However, the behaviors mediated by TFP rely on their dynamics and no quantitative model has been proposed to date to explain how cycles of extension and retraction are controlled. For example, even after decades of research by many groups, fundamental questions like whether there is a molecular ruler that sets TFP length or whether pilus extension/retraction are triggered or stochastic have remained unanswered.

The major hurdles to describing TFP dynamics are the limitations of current approaches for visualizing pili. For example, TFP were first imaged by electron microscopy, but this method can only be performed on fixed or frozen cells such that dynamics are lost^13–15^. Optical tweezers, atomic force microscopy, micropillar assay and traction force microscopy are techniques to measure pilus retraction forces and also yield information about retraction dynamics, but in an indirect way and only for pilus retraction^4,16–21^. A recent study used interferometric imaging to directly image pili in living cells, but this technique generates a strong halo around the cell that overshadows any pili that are shorter than ~3 microns^22^. Despite the limitations of these approaches, they have led to several competing models for how the switch between TFP extension and retraction is controlled. A cryo-EM study did not observe motors at the base of unpiliated structures, suggesting that the motors do not remain bound after TFP retraction^12^. Meanwhile, an interferometry study focusing on the longest subpopulation of TFP suggested that TFP retraction is triggered by surface association^22^. However, the inability to directly visualize the dynamics of the entire TFP population previously limited the ability to directly test these models.

Here we addressed the above limitations by directly fluorescently labeling the TFP of *P. aeruginosa*. Fluorescent labeling of TFP was first achieved with non-specific labeling of extracellular proteins^23^. However, similar to the interferometry approach, this surface labeling approach led to a strong halo from staining of the cell body that prevented analysis of short pili. More recently, TFP from *Caulobacter crescentus* and *Vibrio cholerae* were directly labeled by introducing a reactive cysteine residue into the pilin sequence^24–26^. Here we apply this approach to *P. aeruginosa* and use it to perform the first direct quantitative analysis of full TFP extension and retraction cycles of individual pili. We go on to develop and test quantitative models for the behaviors we observe. We show that TFP production rate, length, and dynamics can be fully explained by the mutually exclusive stochastic binding of the extension and retraction motors, and that this stochasticity persists in the presence or absence of surface association.

## Results

### Quantifying TFP dynamics reveals that P. aeruginosa makes mostly short pili that are highly dynamic

We fluorescently labeled the major protein of the *P. aeruginosa* pilus fiber (PilA) by introducing a cysteine point mutation, A86C, that we then labeled with the thiol-reactive maleimide dye Alexa488-mal (Supplementary Figure 1)^24^. To check that this mutation does not disrupt TFP function, we analyzed twitching motility. Using a standard stab agar twitch assay, we show the PilA-A86C mutant twitches at levels close to wild type on the population level (Supplementary Figure 1). We then looked at individual cells confined between a 0.5% agarose pad and the cover slip and found that cells in this condition twitch actively (Supplementary Movie 1), indicating that the PilA-A86C mutation is functional. We used this configuration for all our experiments unless stated otherwise.

The fluorescent labeling strategy resulted in bright images of dynamic pili with high contrast (Fig. 1a,b and Supplementary Movie 2). Having established that we can label TFP without disrupting their function, we first counted the number of pili that individual cells make in a single snapshot and confirmed previous reports that used electron microscopy to show that only a minority of cells (<25%) are piliated at any given time^15,27^. However, when we then imaged single cells for a period of ~30 seconds, we found that >80% of cells form at least one pilus (Fig. 1c). We quantified the rate of pilus production, *R_p_*, in individual cells and found a very broad distribution between 0 and 35 pili per minute, with a characteristic rate for a typical cell of 8 min^−1^ (Fig. 1d). Whereas static imaging suggested that pili are only made by a small subpopulation of *P. aeruginosa* cells^15,27^, our dynamic imaging suggests that nearly all *P. aeruginosa* cells make short-lived highly-dynamic pili.

**Fig. 1.**
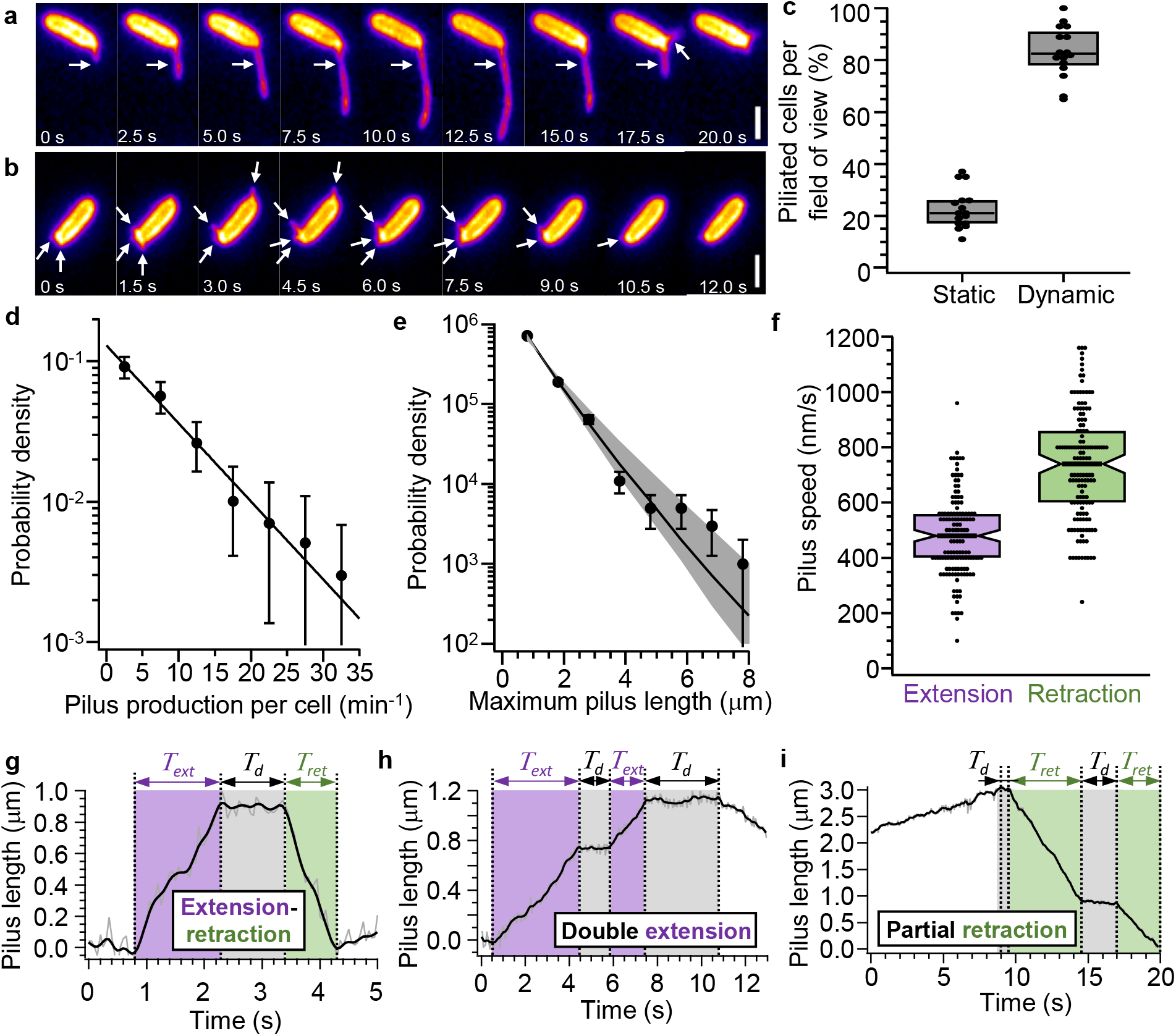
Quantitative measurement of pilus dynamics using Alexa488 coupled to thiolreactive maleimide and the PilA^A86C^ Cysteine knock-in mutant on an agarose pad. **a**, Movie frames showing the extension and retraction of a long pilus (white arrow, *L_p_* = 5 μm). Scale bar 2 μm. **b**, Movie frames showing the typical extension and retraction of several short pili (white arrows, *L_p_* ≤ 1 μm). Scale bar 2 μm. **c**, Comparison of fraction of cells in a single image with at least one pilus when analyzed in just a single frame (static) or a movie (dynamic) of 30 seconds in length. **d**, Distribution of pilus production rate per cell. Error bars are 95% confidence intervals obtained by bootstrapping. **e**, Distribution of maximum extension lengths of individual pili. Error bars are 95% confidence intervals obtained by bootstrapping. Gray shaded area: 95 % confidence interval from model simulation for comparison (MCS, see below and Methods). **f**, Pilus extension and retraction speed. Notched boxes represent the median and 25 % / 75 % quantiles. **g**, Time trace of pilus length for a typical pilus extension/retraction event: extension time *T_ext_*, dwell time *T_d_*, retraction time *T_ret_*. **h**, Time trace of pilus length for a double extension event. **i**, Time trace of pilus length for a discontinuous retraction event. **g,h,i** Also see Supplementary Figure 2 and Supplementary Movies 3,4,5. (See Supplementary Table 4 for sample sizes and number of replicates).

To further quantify TFP behavior we measured the distribution of pilus lengths (Fig. 1e). We found that the pilus length (*L_p_*) also exhibits a wide distribution between 0.3 μm (limited to optical resolution) and 8 μm, with a characteristic length for a typical pilus of 0.8 μm. We note that this result differs from the only other quantitative study of *Pseudomonas* pilus lengths which observed only pili longer than 3 μm^22^. However, the interferometric imaging technique used in that study could not detect pili shorter than the halo produced by the cell itself (2 – 3 μm). Our direct labeling approach supports the hypothesis that most *P. aeruginosa* extend short, short-lived pili.

### TFP extension and retraction events can be discontinuous

The ability to directly label pili enabled us to analyze the extension and retraction dynamics of individual pili. A typical pilus had a median extension speed of *v_ext_* = 500 nm/s and a median retraction speed *v_ret_* = 750 nm/s (Fig. 1f). These rates are in agreement with a previous study that measured extension and retraction velocities in *P. aeruginosa*^23^, further validating that our measurements reflect the physiologically-relevant behaviors of *P. aeruginosa* TFP.

An analysis of the durations of extension and retraction proved more surprising. We analyzed the entire extension-retraction cycle by tracing the tips of pili relative to the cell body over time and defining periods of extension, dwelling, and retraction (see Methods and Supplementary Figure 2). A typical pilus extends for about *T_ext_* = 2 seconds, then dwells for less than *T_d_* = 1 second, and finally retracts all the way back (Fig. 1g and Supplementary Movie 3). We also observed unexpected patterns of extension and retraction. In 15 out of 196 dwell events, an extension event was followed by another extension (Fig. 1h and Supplementary Movie 4). Similarly, in 11 out of 127 retraction events, the pilus stalled during the retraction, resulting in another dwell event, followed by continued retraction as shown in Fig. 1i and Supplementary Movie 5. Such intermittent dwell events resulting in discontinuous extension and retraction represent a previously unappreciated feature that must be explained by any model of pilus dynamics.

### TFP extension and retraction dynamics are unaffected by the presence of a surface

To understand the mechanisms that control pilus dynamics and the biophysical basis for our surprising findings we next sought to understand how the switch between TFP extension and retraction is coordinated. A prominent hypothesis is that TFP retraction is triggered by mechanical contact of the pilus tip with a surface^12,22^. To test this model, we compared the TFP dynamics of cells in two different conditions: cells confined between agarose and a coverslip (surface-associated), and cells prevented from contacting a surface by holding them 5 μm above the cover slip using an optical trap (liquid-trapped) (Fig. 2a). In addition to holding the bacteria away from the surface, the line-scanning optical trap^28^ orients the cells with the microscope focal plane, which allowed us to observe pilus dynamics on both cell poles.

**Fig. 2.**
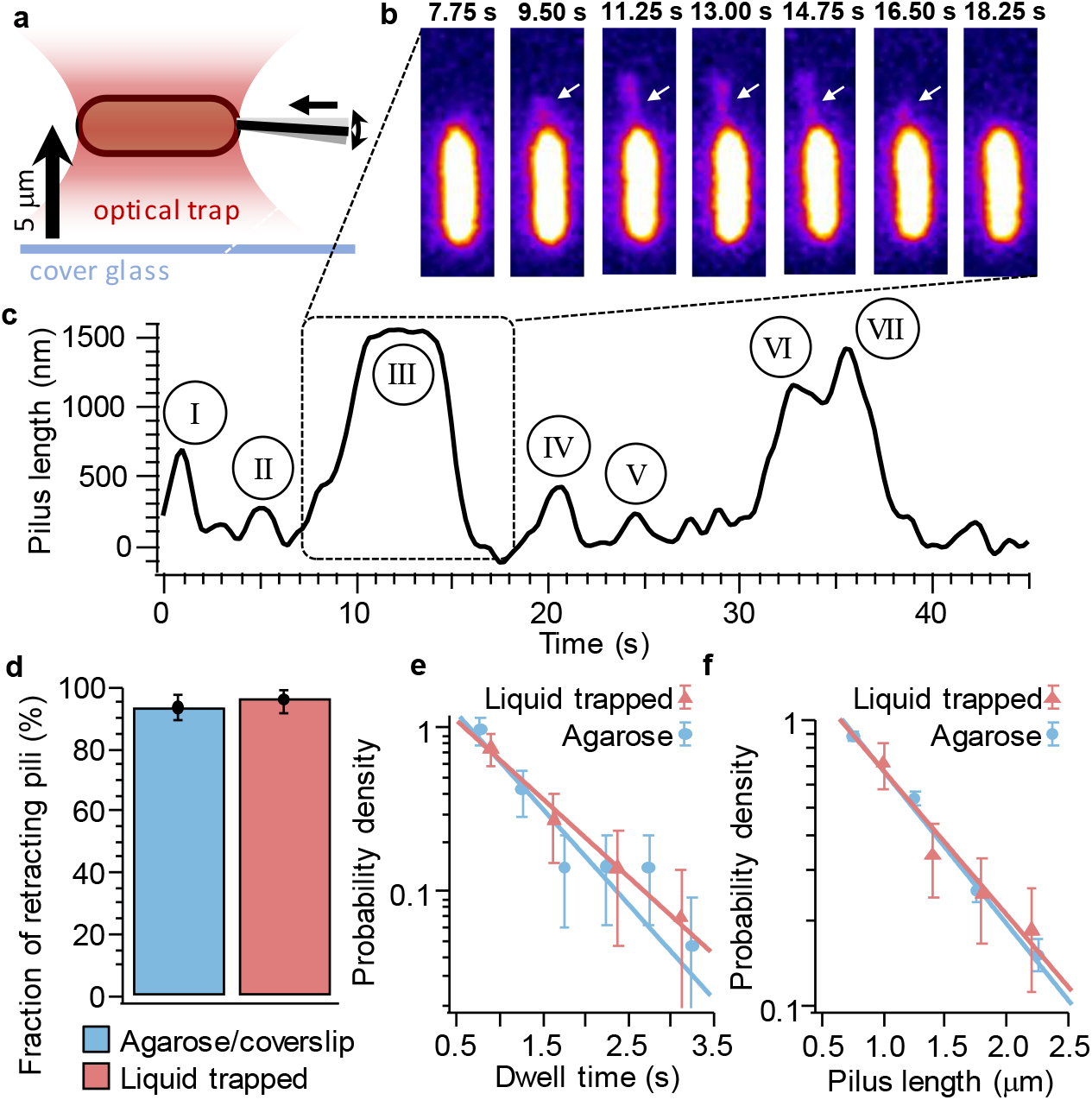
Pilus retraction does not require mechanical stimulation. **A**, Schematic of surface contact-free (“liquid-trapped”) assay: single cells are held about 5 μm above the surface and aligned with the focal plane by line-scanning optical tweezers. **b**, Image sequence of an individual pilus extending and retracting without surface contact (also see Supplementary Movie 10). **c**, Time trace of pilus length for seven individual pili (roman numerals) extending and retracting from the same pole of the same cell without surface contact. **d**, Fraction of retracting pili for cells with and without surface contact. **e**, Dwell times between stop of extension and start of retraction of individual pili for cells with and without surface contact. **f**, Maximum length of individual pili for cells with and without surface contact. (See Supplementary Table 4 for sample sizes and number of replicates).

As show in Supplementary Movie 6 and 7, individual cells with labeled pili confined between 0.5% agarose and the coverslip can twitch, which means that their TFP are in mechanical contact with the environment. Similarly, we observed frequent TFP extension and retraction for liquid-trapped cells (Fig. 2b,c and Supplementary Movies 8-10), indicating that loss of surface contact does not completely abolish pilus retraction. If the surface-triggered model is true, the dynamics of TFP for cells with and without surface contact should be quantitatively different. For example, if mechanical contact of the pilus tip with a surface triggers pilus retraction, then we expect to see fewer retracting TFP for liquid-trapped cells compared to surface-associated cells. Surprisingly, the fractions of retracting pili are 95% for liquid-trapped cells and 93% for surface-associated cells and are therefore indistinguishable (Fig. 2d). It is possible that we did not observe a difference in the fraction of retracting pili because surface-association accelerates the timing between extension and retraction, in which case TFP would retract eventually even without a mechanical trigger signal. To test this hypothesis, we quantified the time each TFP dwells between when the extension comes to a halt and the retraction starts. If the contact with a surface stimulates pilus retraction the distribution of dwells times should be shorter for surface-associated cells. However, the distributions of dwell times for both conditions were indistinguishable from each other (Fig. 2e). Similarly, the distributions of TFP length were indistinguishable in both conditions (Fig. 2f), indicating that surface contact also does not stop TFP extension. We therefore conclude that the dynamics of the switch between pilus extension and retraction are indistinguishable whether or not a surface is present.

### A stochastic model of motor binding can explain the full extension and retraction cycles of individual TFP

After the surprising result that TFP dynamic are unaffected by the presence of a surface we considered other mechanisms that could explain the switch between extension and retraction. A recent cryo-EM study suggested that only one type of motor, extension or retraction, is bound to the pilus machine at any given time^12^. This indicates that both motors must compete for the binding to the machine. Furthermore, the distributions of the maximum pilus length, the rate of pilus production, and the dwell time between extension and retraction are exponential in shape (Fig. 1d,e and Fig. 2e), suggesting that stochastic protein binding and unbinding might govern pilus dynamics (see Methods). We thus formulated a quantitative model in which pilus extension and retraction are governed by the stochastic binding of an antagonistic extension or a retraction motor to the pilus base in a mutually exclusive manner. We note that the only assumptions of this model are that each motor has a finite probability to bind the unbound pilus machine, and that no more than one motor can be bound at a given time (Fig. 3a).

**Fig. 3.**
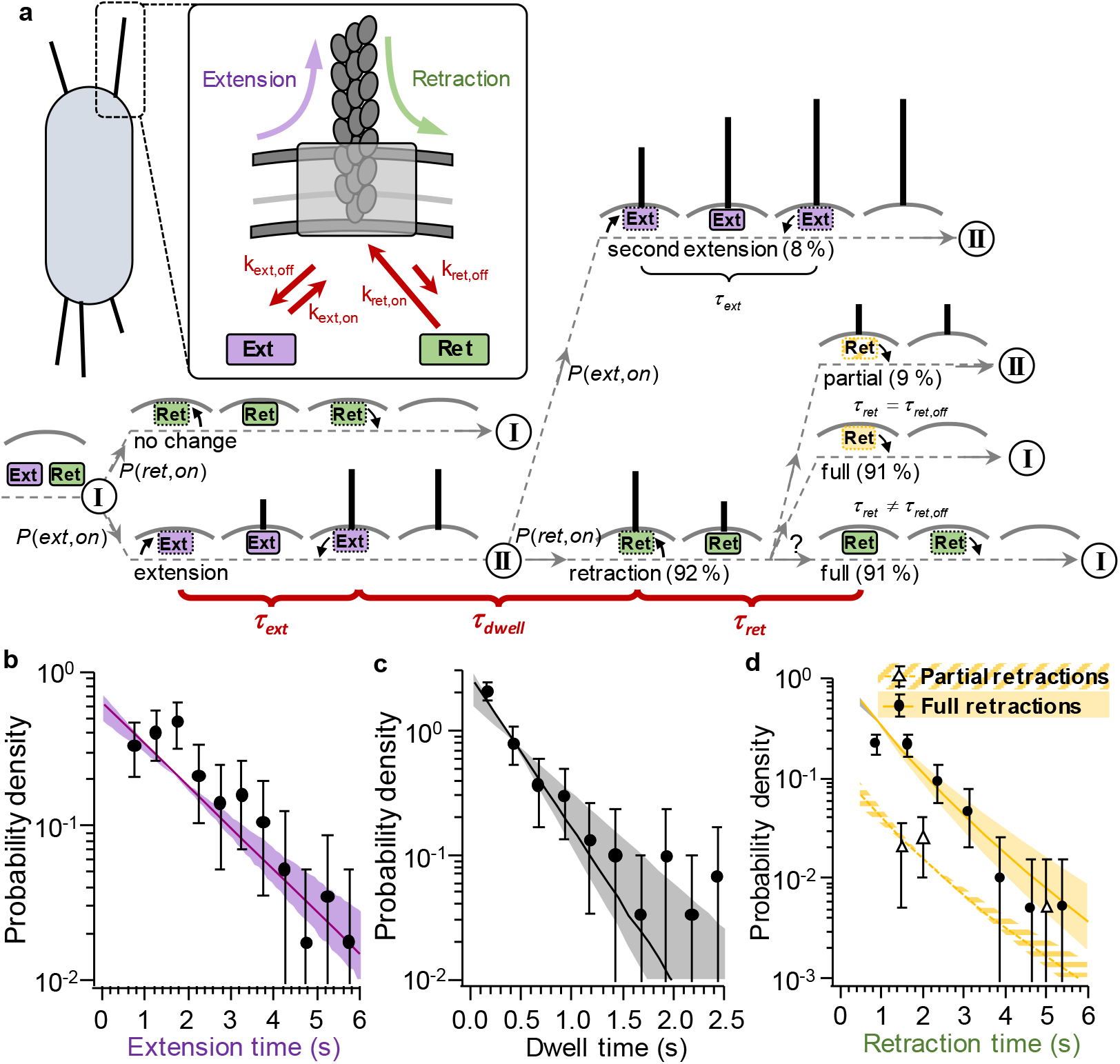
Competitive substrate binding model predicts rare multistep extension and retraction events with short intervening stalls. **a**, Model schematic. The extension motor (Ext, purple) and retraction motor (Ret, green) bind with probability *P*(*Ext, on*) and *P*(*Ret, on*), respectively. I and II denote, respectively, un-piliated and piliated pilus machine without bound extension or retraction motor. **b**, Histogram of extension times of individual pili. **c**, Histogram of dwell times between stop of pilus extension and start of the subsequent pilus retraction. **d**, Histogram of retraction times of individual pili. **b, c, d** Error bars are 95 % confidence intervals obtained by bootstrapping. Shaded areas are 95 % confidence interval from model simulations (MCS, see Methods). (See Supplementary Table 4 for sample sizes and number of replicates).

This stochastic model for TFP dynamics includes six independent parameters: the extension and retraction speed of the pili (*v_ext_* and *v_ret_*), the binding and unbinding rates of the extension motors (*k_ext,on_* and *k_ext,off_*), and the binding and unbinding rates of the retraction motors (*k_ret,on_* and *k_ret,off_*). The extension and retraction speeds were directly measured (Fig. 1f). In the following, we show how each of the other rates can be estimated from our data (see Fig. 3 and Methods for details). We then use our model to make quantitiative preditions that we validate experimenally and show how the model makes the unexpected prediction that the main limiting factor for pilus dynamics is the unbinding of the retraction motor.

The duration of each pilus extension event is equal to how quickly the extension motor becomes unbound. Thus, the unbinding rate of the extension motor can be derived from the characteristic unbinding time *τ_ext_* by 1/*k_ext,off_* = *τ_ext,off_* = *τ_ext_*. We directly measured the distribution of pilus extension times (Fig. 3b), which had an exponential shape and a characteristic time of 1.6s (*τ_ext_*, Fig. 3b), indicating that 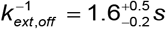.

The relationship between the unbinding rate of the retraction motor and the duration of retraction events is more complicated. For the majority of retraction events, the pilus becomes fully retracted so we cannot tell when the retraction motor becomes unbound. We do, however, observe a number of retraction events that are interrupted by a dwell period, suggesting that the retraction motor became unbound during these events. We observe such events with a probability of 11 partial retractions out of 127 total events (9%). These events represent the short-time tail of the distribution of unbinding times. To account for all retraction events, we used a maximum likelihood approach to find the characteristic time constant of unbinding that best accounts for the full distribution of both complete and partial retraction events. We note that the only assumption in this approach is that the retraction unbinding times are exponentially distributed, which is consistent with all our other pilus measurements. As detailed in the Methods and Supplementary Figure 3, this maximum likelihood approach estimated the unbinding rate of the retraction motor as 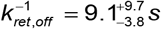.

The dwell periods *T_d_* that follow every extension event allow us to estimate the binding rates of both extension and retraction motors. The time to the next extension or retraction event is set by the binding of the next motor to that pilus, such that *τ_dwell_* = 1/(*k_ext,on_* + *k_ret,on_*). We measured a characteristic dwell time of 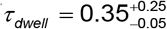 s from the distribution of all dwells (Fig. 3c). The ratio of the binding rates of the extension and retraction motors sets the fraction of post-dwell events that are extensions versus retractions. As described above, post-dwell we observed 15 secondary extensions and 181 retractions, suggesting *k_ext,on_*/*k_ret,on_* = 15 / 181. Combining these values and taking into account the finite experimental time resolution that limits our ability to detect short dwell periods (see Methods and Supplementary Figure 4), we estimate 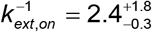 s and 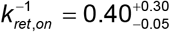 s.

To validate our model and parameters, we sought to use the model to predict the ratio of partial to full retractions. We simulated cycles of extension and retraction of individual pili using the Monte Carlo method by drawing random samples from the model’s distributions of extension and retraction times and velocities (referred to as MCS, see Methods). From those numbers, we calculated the expected length of each pilus and determined if its retraction time was enough to fully retract it, i.e., if the retraction gave rise to a partial or full retraction. We compared our simulated distribution of partial retractions (Fig. 3d, yellow dashed) and full retractions (Fig. 3d, yellow) to our experimental findings and found good agreement. As a further verification, we analyzed an independent set of data that was not used to estimate the model’s parameters and found that the resulting pilus lengths agreed well with our model’s simulated results (Fig. 1e).

### The effect of a retraction motor mutant on discontinuous retractions is accurately predicted by the stochastic TFP model

To further support our model we used a genetic approach to test one of its predictions. The model suggested that if TFP extension and retraction speeds are independent of motor binding rates, a mutant that reduces retraction speed should show more partial retraction events because TFP need more time to complete a full retraction. To test this prediction we analyzed pilus dynamics in a point mutant (PilT-H222A) in the ATPase activity of the PilT retraction motor that affects pilus retraction speed^29^. We first confirmed that PilT-H222A pili retract three times slower compared to WT, while the pilus extension speed and all four binding/unbinding rates remain indistinguishable from WT (Supplementary Figure 3,5). We then measured the fraction of partial retractions of PilT-H222A pili, and indeed found that they increased relative to WT (Fig. 4a). We also performed a simulation in which we reduced *v_ret_* threefold but left all the other parameters unchanged and observed good agreement between this simulation and our experimental results with PilT-H222A (Fig. 4b). These results show that the discontinuous pilus retraction can be explained quantitatively by the stochastic binding and unbinding of the pilus motors.

**Fig. 4.**
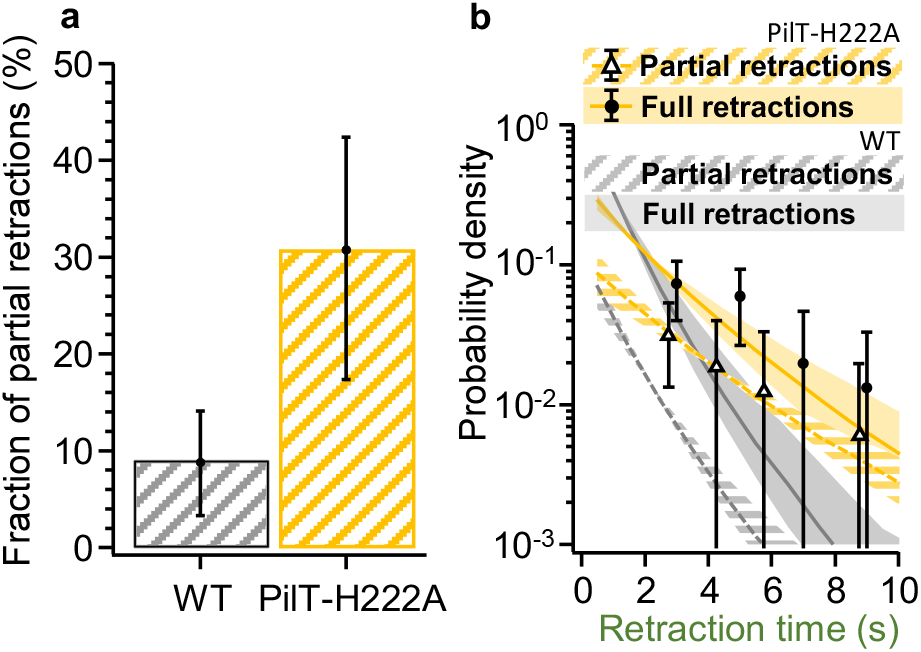
The change of the fraction of partial retractions for the slowly retracting mutant PilT-H222A is accurately predicted by the model. **a**, The fraction of partial retractions increases about threefold from WT to PilT-H222A. **b**, Distribution of retraction times of individual pili for PilT-H222A (yellow = model prediction, markers = experimental data) and WT (grey) for comparison. **a, b**, Error bars are 95 % confidence intervals obtained by bootstrapping. **b**, Shaded areas are 95 % confidence intervals from model simulation (MCS, see Methods). (See Supplementary Table 4 for sample sizes and number of replicates).

### The switch between extension and retraction is governed by the stochastic binding and unbinding of both motors

We next sought to use our quantitative framework to determine how TFP switch between extension and retraction. Based on our observations and recent cryo-EM and interferometric imaging data we tested three competing models^12,22^: One hypothesis is that both the binding and unbinding of the retraction motor are purely stochastic and independent of the presence of the pilus itself (Model 1: The stochastic model). A second possibility is that the retraction motor can only bind to the machine if a pilus is present, but unbinds in a stochastic manner whether or not the pilus has fully retracted (Model 2: The pilus-dependent model). A third possibility is that the retraction motor both only binds if a pilus is present and unbinds as soon as the pilus is fully retracted (Model 3: The pilus-sensing model). These three models make different predictions for the TFP production rate. Due to the rapid on rate and slow dissociation rate of the retraction motor compared to the extension motor, Model 1 predicts that the pilus machine is occupied by the retraction motor most of the time. Because the extension and retraction motors compete for binding to the pilus machinery, this suggests that the rate of pilus production is primarily limited by the retraction motor. In Models 2 and 3 the retraction motor does not bind the unpiliated machine, and thus the extension motor can bind more frequently, resulting in more pilus extension events compared to Model 1. Furthermore, since the retraction motor unbinds after the pilus is fully retracted in Model 3, we would expect to see the largest number of pili in this model.

To differentiate between these different behaviors of the retraction motor, we again used the Monte Carlo method (see Methods) to simulate cycles of the stochastic binding and unbinding of the extension and retraction motors using each of the three models (Fig. 5a). We counted the number of pilus extension events per pilus machine in a 60 second time window in the simulation and found that for the simple stochastic model (Model 1), the pilus production rate was approximately exponentially distributed with typically one pilus event per minute (Fig. 5c). The simulations for Models 2 and 3 were distinctively different from those of Model 1 as both Models 2 and 3 displayed a more Gaussian distribution peaking between 3 pili per minute (Model 2) and 6 pili per minute (Model 3).

**Fig. 5.**
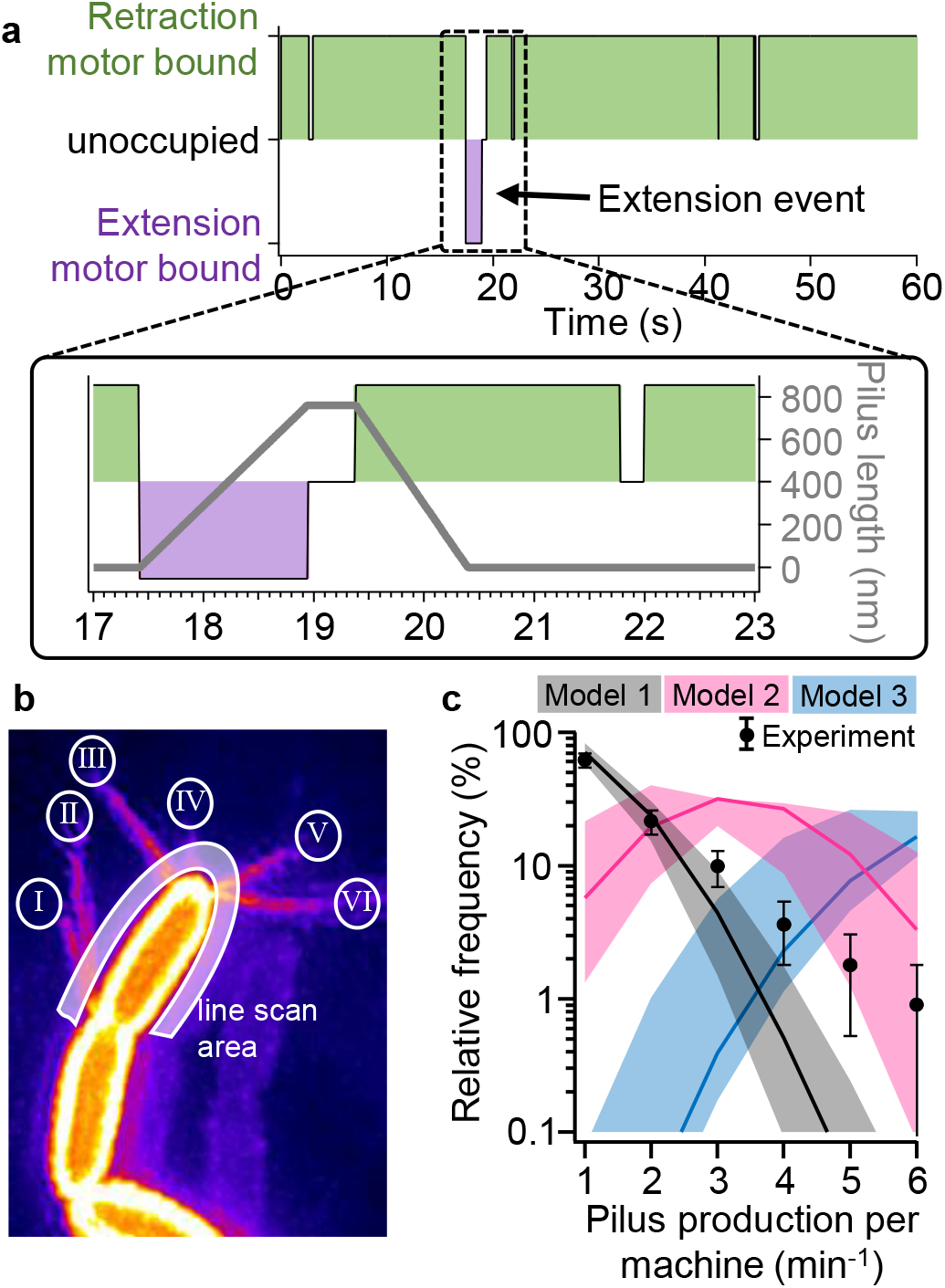
Comparison of the pilus production rate predicted by different models for the switch between extension and restriction. **a**, Example of Monte Carlo simulation for binding and unbinding of the extension and retraction motor showing a single pilus extension event. Note that the retraction motor stays attached after the pilus is retracted fully. **b**, Maximum projection of 60 super-resolved movie frames showing directions of all pili that have been extended by the cell. Roman numerals label individual pilus machines. Thick transparent curve represents the line scan area used to analyze pilus production see (Supplementary Information). **c**, Distribution of the pilus production rate per machine. Experimental data are shown as black dots with error bars. Monte Carlo simulations are shown for the stochastic model (Model 1, grey), the pilus sensing model (Model 2, pink), and the pilus-dependent model (Model 3, blue). Error bars are 95 % confidence intervals obtained by bootstrapping. Shaded areas are 95 % confidence intervals from model simulation (MCS, see Methods) and bold lines are their means. (See Supplementary Table 4 for sample sizes and number of replicates).

To experimentally differentiate the three hypotheses, we measured the pilus production rates of individual pilus machines and compared these results to the simulated distributions from the three models. Measuring the pilus production rate of individual machines is experimentally challenging because a pilus machine is only 15 to 20 nm in diameter and neighboring complexes can be closer together than the conventional optical resolution limit^12^. To tackle this problem, we used live-cell super-resolution microscopy and looked at maximum projections of entire movie stacks (Fig. 5b and Supplementary Movie 11). Due to the strong curvature at the poles, pili originating from close-by machines (roman numerals in Fig. 5b) emanate at different angles and can be more easily distinguished. We thus analyzed changes in intensity along a line just outside the cell circumference (transparent curve, Fig. 5b) in a kymograph (Supplementary Figure 6b). By assigning each pilus extension event to the machine from which it emanated, we were able to count the frequency of pilus extension events per individual machine (Supplementary Figure 6c). We found that pilus extension frequency was exponentially distributed with an average of roughly one pilus extension events per minute. Qualitatively, these data agreed well with the simple stochastic model (Model 1), but were incompatible with both Models 2 and 3 since these distribution have a different shape (Fig. 5c).

The only deviation between our Model 1 simulation and the experimental results is for production rates ≥ 4 pili min^−1^. We suggest that this small deviation can be attributed to our finite imaging resolution. This resolution limit makes it difficult to distinguish if two short pili emanate from the same machine or from two nearby complexes, which in turn leads to a systematic overestimation of pilus extension frequency. Nevertheless, we quantitatively tested if this deviation is statistically significant and performed Kolmogorov-Smirnov tests that compare the simulations for each model using the mean binding/unbinding rates and their lower and upper bounds separately to the experimental data. Model 1 has P < 0.05 for the mean binding/unbinding rates, P < 0.001 for the lower bound of these rates and P > 0.05 for their upper bounds. Models 2 and 3 yield P > 0.05 for all combinations of the rates. This confirms the qualitative result that Model 1 can best explain the experimental data and further indicates that our estimates for the binding and unbinding rates of the motors might be slightly higher than the actual binding and unbinding rates. Together, our findings support the conclusion that the switch between pilus extension and retraction is stochastic and that the rate of pilus production is limited by the slow unbinding step of the retraction motor.

## Discussion

Here we fluorescently labeled the TFP of *P. aeruginosa* and quantified the extension and retraction cycles of individual pili. We confirmed previous findings of pilus extension and retraction rates and the number of piliated cells at any given time. However, we also made the surprising finding that cells make many more and much shorter pili than previously appreciated. For example, while prior studies have focused on long pili (> 3 μm) that can be detected by interferometry, we showed that pili are predominantly shorter than 1 μm^22,23^. Further, we were able to show that pili are highly dynamic: a typical cell makes a new pilus every 5 – 10 seconds and retracts each pilus rapidly. Moreover, both extension and retraction can be discontinuous. By quantitatively comparing different models for the switch between extension and retraction we were able to show that only the stochastic binding and unbinding of extension and retraction motors is able to quantitatively explain all TFP dynamics, and that these behaviors are not altered by the presence of a surface. Below we discuss the molecular, biophysical, and physiological implications of these findings.

The model we propose here is minimal in its nature and relies only on the presence of extension and retraction motors and their stochastic interactions with the pilus machine. We suggest that cells can tune the binding and unbinding rates that we present here to alter pilus dynamics. An interesting conclusion from our results is that the major throttle of pilus extension is the low unbinding rate of the retraction motor. In *Myxococcus xanthus*, only one pole is piliated leading to directed twitching motility. In agreement with our model, retraction motors localize predominately to the lagging pole that does not make TFP while extension motors localize to the piliated leading pole^30^. In contrast, our finding that the retraction motors likely remain bound to the pilus machines well after pilus retraction is complete does not agree with the interpretation of a cryo-EM study of *M. xanthus* in which the retraction motor’s electron density was not seen bound to unpiliated machines^12^. This discrepancy could be due to an artifact introduced during the interpretation of the cryo EM pictures or during sample preparation. Another possibility is that TFP dynamics differ across species. Future structural studies in *P. aeruginosa* and TFP dynamics measurements in *M. xanthus* should help understand these differences.

While our results indicate that TFP dynamics can be understood as a simple competitive binding system, this system can still be regulated by accessory factors that alter the base rates of binding, unbinding, extension, and retraction. For example, in *M. xanthus*, the direction of twitching motility and the TFP localization pattern oscillates periodically and this process is orchestrated by the frz system, suggesting that frz regulates both motors either directly or indirectly^31^. Similarly, in *P. aeruginosa*, the Pil-Chp two-component system regulates pilus behaviors, both through biochemical interaction of the two response regulators PilG and PilH with the pilus machine and by transcriptional modification using the cAMP dependent transcriptional regulator Vfr^32–34^. The second messenger c-di-GMP has also been shown to interact directly with the extension motor and other components of the TFP machine^35–37^, thus resembling another interesting candidate for the regulation of the binding and unbind rates of TFP.

Previous studies have invoked complicated molecular force sensors and fast coordination between motor elements. However, our findings indicate that such elaborations are not necessary to explain basic pilus behaviors. Nevertheless, our results do not exclude the possibility that surface contact is sensed actively by pilus retraction leading to subsequent biochemical signaling that changes protein synthesis or transcription^5,25,32^. While the presence of a surface does not alter TFP dynamics directly, surface sensing could still be mediated by TFP through biochemical changes in the pilus machine that do not affect the binding or unbinding rates but are sensed by auxiliary modules like PilJ.

We also note that our model abstracts the extension and retraction motors. In *P. aeruginosa*, PilB is the only known extension motor, PilT is considered the primary retraction motor, and PilU has been shown to affect retraction^22,38^. Our analysis confirms that PilU is indeed not needed for retraction but does affect retraction speed (Supplementary Figure 6). Recent studies also suggest that these motors may have more complicated interactions^38,39^, and in the future our model could help tease apart the specific contributions of different mutants to the extension and retraction cycle.

Our findings show that while most *P. aeruginosa* make many pili, these cells have tuned the affinities and rates of the extension and retraction motors to generate short pili that are rapidly and fully retracted. However, this also prevents individual pilus machines from rapidly extending new pili after a retraction event, such that frequent pilus extension requires the presence of multiple pilus machines. We suggest that tuning the pilus parameters to increase retraction events benefits *P. aeruginosa* by enhancing surface interactions such as the displacement required for twitching motility. Frequent pilus retraction also allows planktonic cells to efficiently sample the environment for the presence of a surface. Once a pilus is bound and retracts under load, subsequent downstream signaling activates transcriptional programs associated with a surface bound lifestyle^5,11,25,32,40^. Similarly, tuning the parameters to ensure that most pili are fully retracted enhances pilus subunit recycling to the membrane, thereby enhancing the rate of new pilus production. Thus, our findings support the hypothesis that *P. aeruginosa* has evolved to maximize its pilus budget for interaction with surfaces. In the future it will be interesting to see how regulatory elements such as the Pil-Chp two-component system or second messenger mediated modifications can alter the base rates described here^32,35-37^. Further, it is likely that other species with other physiological demands and constraints modulate the kinetics of motor binding to change pilus length, number, and dynamics to achieve other functions, like cell-cell interactions or DNA uptake.

## Methods

### Strains, plasmids, growth conditions, and cloning

The bacterial strains, plasmids, and primers used in this study are described in Supplementary Tables 1 – 3.

*P. aeruginosa* PAO1 was grown in liquid LB Miller (Difco) and Cysteine free EZ rich defined medium (Teknova)^41^ in a floor shaker, on LB Miller agar (1.5 % Bacto Agar), on Vogel-Bonner minimal medium agar (VBMM, 200 mg/l MgSO_4_ 7H2O, 2 g/l citric acid, 10 g/l K_2_HPO_4_, 3.5 g/l NaNH_4_HPO_4_ 4 H_2_O, 1.5 % agar), and on no salt LB agar (NSLB, 10 g/l tryptone, 5 g/l yeast extract, 1.5 % agar) at 30 °C (for cloning, see below) or at 37 °C. *E. Coli* S17 was grown in liquid LB Miller (Difco) in a floor shaker and on LB Miller agar (1.5 % Bacto Agar) at 30 °C (for cloning, see below) or at 37 °C. Antibiotics were used at the following concentrations: 200 μg/ml carbenicillin in liquid (300 μg/ml on plates) or 10 μg/ml gentamycin in liquid (30 μg/ml on plates) or 10 μg/ml anhydrotetracycline in liquid for Pseudomonas and 100 μg/ml carbenicillin in liquid (100 μg/ml on plates) or 30 μg/ml gentamycin in liquid (30 μg/ml on plates) for E. Coli.

#### The *ΔfliC* deletion and PilA-Cysteine knock-in strains

were generated using two-step allelic exchange^42^. Briefly, the Δ*fliC* cloning vewere concentrated intoctor was created by digesting the pEXG2 backbone with the HindIII HF restriction enzyme (NEB). 500 bp of the flanking regions up- and downstream of *fliC* were PCR amplified using primer pairs DfliC_P1/2 and Dflic_P3/4. Both products were then joined using sewing PCR with primers Dflic1/4 and subsequently digested with HindIII HF. The product was then ligated into the pEXG2 backbone using T4 DNA ligase (NEB). The PilA-Cysteine knock-in vectors were created similarly by amplifying the 500 bp flaking regions up- and downstream of the mutation site using primers pilA-XYYC, where XYY stands for the name and location of the original residue. The overlapping primers were chosen as reverse complement containing the point mutation. After ligation, the cloning vectors were electroporated into *E. Coli* and the correct mutation was confirmed using PCR and sanger sequencing with primers pEXG2_Ver1/2. For mating, 1.5 ml *E. coli* containing the vector were grown to OD 0.5. The *P. aeruginosa* parental strain was grown overnight, and 0.5 ml culture was diluted 1:2 into fresh LB and incubated for 3 hours at 42 °C. Both cultures were concentrated into 100 μl and spotted onto an LB agar plate and incubated overnight at 30 °C. The puddle was scrapped off, resuspended into 150 μl PBS, spread onto a VBMM plate containing 30 μg/m1 gentamycin and incubated 24 hours at 37 °C. Six single colonies from the VBMM plate were struck onto NSLB and incubated for 24 hours at 30 °C. Several single colonies from the NSLB plate were screened for the correct mutation using PCR amplification with the flaking primers and sanger sequencing.

#### The chromosomal tetracycline (tet) inducible *pilT-H222A* mutant

was constructed in two steps: first, the plasmid pMK47 containing an inducible mKate2 construct was constructed by digesting the pUC18-mini-Tn7T-LAC vector with NsiI and Eco53kI restriction enzymes (NEB)^43^. A tet regulation cassette was PCR amplified from plasmid pXB300^44^ using primers pMK47_F1.For and pMK47_F1.Rev and the gene coding for the fluorescent protein mKate2 was amplified from plasmid pPaQa^5^ using primers pMK47_F2.For and pMK47_F2.Rev. All three fragments were joined using Gibson assembly. Next, plasmid pMK73 was made by PCR amplifying the backbone of plasmid pMK47 containing the tet inducible construct using primers pMK47BB.For and pMK47BB.Rev. The PilT-H222A fragment was generated by a two-step PCR amplification: first, the regions up-downstream of the mutated residue were amplified using primer pairs pMK73F1.For / PilT_H222A_P2 and PilT_H222A_P3/pMK73_Flag, introducing an additional Flag tag at the C-terminus of PilT. The overlapping primers were designed as reverse complements containing the point mutation. Then, both fragments were joined using sewing PCR with primers pMK73F1.For and pMK73F1.Rev. This fragment and the backbone were then joined using Gibson assembly. Plasmid pMK47 and pMK73 were inserted into the chromosome of PAO1 by coelectroporation with plasmid pTNS2^43^. In brief, 10 ml of the parental strain was grown to late log phase (OD 1.0), washed three times and then resuspended in 60 μl 300 mM sucrose together with 600 ng of pTNS2 and 600 ng of either pMK47 or pMK73. After electroporation, strains were recovered in 1 ml LB for 2 hours at 37 C shaking and the entire reaction was plated onto LB agar containing 30 μm/ml gentamycin. Single colonies were verified using sanger sequencing.

### Sample preparation and imaging

For imaging of pilus dynamics, cells were grown overnight in EZ rich medium at 37 °C, diluted 1:1000 into fresh EZ rich and grown to mid log phase (OD 0.4). EZ rich medium has a low background fluorescence and the absence of free Cysteine improves the labeling efficiency with the maleimide dye while assuring rich growth conditions. 1 mg of Alexa488 maleimide (Fisher A10254) was suspended in 400 μl DMSO, aliquoted and stored at −20 °C. Freeze-thaw cycles were avoided as they degrade efficiency of pilus labeling. Dye was added 1:100 to 180 μl of culture and incubated for 45 minutes at 37 °C in the dark. Cells were washed twice gently in EZ rich by pelleting at 6 krpm for 30 seconds in a conventional tabletop centrifuge and concentrated to 20 μl. For optical trapping experiments in liquid, a tunnel slide was made by placing a regular cover slip on a microscope slide, separated by double sided sticky tape at each side of the cover slip. Cell were flushed in by capillary forces using a pipette and ends were sealed with Valap to prevent evaporation and flow of liquid. WT cells have flagella and typically swim out of the optical trap. To prevent cells from leaving the trap, we used a flagella knockout Δ*fliC* for all quantitative experiment after confirming qualitatively that flagellated WT cells still make and retract pili when trapped. For all other experiments, 0.5 % agarose pads were made by melting 1.0 % agarose in water. Agarose was cooled down to 60 °C and mixed 50:50 with double concentrated EZ rich at 60 °C. 1 ml of labeled cell culture was spotted on each pad and the pad was transferred to a no 1.5 glass bottom petri dish (Mattek). All experiments were performed at 37 °C on three different microscopes as described in the following.

#### HILO

Highly inclined thin illumination microscopy (HILO) is a variation of total internal reflection microscopy (TIRF)^45^. Similar to TIRF, HILO has a significantly improved signal to background ratio compared to epifluorescence but maintains the axial penetration depth of epifluorescence, thus enabling to observe processes away from the coverslip surface at much reduced bleaching and improved image quality. We used a commercial Nikon Ti-E microscope equipped with a TIRF module and set the direction of the incident laser slightly below the critical TIRF angle. The microscope was used with perfect focus, a 100x NA 1.49 Apo TIRF lens (Nikon), an EMCCD camera (iXon Ultra DU-897U, Andor), a stage top incubator (INU, Tokai Hit) and controlled by Nikon Elements software.

#### R-HILO and optical trapping

The combined optical trapping (OT) and ring-HILO (R-HILO) setup was custom built on a TE2000 body (Nikon). Similar to ring-TIRF^46^, ring-HILO improves the spatial homogeneity of the image by reducing the spatial coherence of the illumination. To our knowledge, this is the first time that HILO has been used in a scanning configuration. Briefly, a 5 W 1064 nm laser (Spectra Physics) was focused in the focal plane using a 100x NA 1.49 Apo TIRF lens (Nikon) to create an optical trapping potential^47^. To form the line optical trap^28,48^, the laser focus was scanned at 200 Hz over a distance of 5 μm in the focal plane of the objective lens using a tip-tilt piezo mirror (Mad City Labs) and a function generator (GFG-8215A, Instek). The laser power was set as low as possible to avoid photodamage of the trapped cell. Samples were positioned using a three-axis piezo stage (Mad City Labs). For R-HILO, a Coherent Obis 488 nm LS 60 mW laser was focused in the back focal plane of the objective lens and scanned on a ring in the back focal plane using a two-axis galvanometric scanner (GVS212, Thorlabs) positioned in a conjugate plane. The radius of the focus with respect to the optical axis and the scanning frequency were set using a NI PCIe-6251 digital to analog DAQ card (National Instruments). Fluorescence excitation, emission, and the optical trapping light were combined using a quad-band dichroic mirror (Di01-R405/488/561-25×36, Semrock), quad-band emission filter (FF01-446/523/600/677-25, Semrock), and a short pass filter (FESH0750, Thorlabs). Images were acquired using a water cooled EMCCD camera (iXon Ultra DU-897, Andor). The entire microscope was controlled using custom written software in National Instruments LabView. Cells were incubated using a custom built, laser-cut incubation chamber and a PID temperature controller (In Vivo Scientific).

#### SIM

We used a Nikon Ti-E N-SIM microscope equipped with perfect focus, a 100x NA 1.49 Apo TIRF lens (Nikon), an EMCCD camera (iXon 3 DU-897E, Andor), a stage top incubator (INU, Tokai Hit) and controlled by Nikon Elements software for structured illumination microscopy (SIM)^49^. Nine images in 2D mode were acquired for every super-resolved image to ensure a high effective frame rate (1 Hz).

### Image analysis and pilus tracking

Images were analyzed in Fiji Briefly, stacks of individual cells were cropped, interpolated 10x to improve further image processing (see below), and photobleaching was removed using the bleach correction tool. A line ROI was drawn along the pilus extending the maximum length of the pilus by at least 1 μm, and the intensity along the ROI was measured for every image of the stack using a macro^50^. These data were than copy and pasted into another software (Wavemetrics Igor Pro) for subsequent processing with custom written scripts. The length of the pilus was detected in every frame using thresholding of the intensity. The resulting kymographs and pilus length trajectories are shown in Supplementary Figure 2. The extension, dwell, and retraction times, pilus length, extension and retraction velocities were extracted from these trajectories semi-automated by identifying start and stop of each extension and retraction by hand.

### Statistical analysis

The number of independent replicates and analyzed samples used in each figure is shown in Supplementary Table 4. As shown in Supplementary Table 4, the parameters of the model were estimated based on data (Fig. 2) that were independent of the data shown in the rest of the paper (Figs. 1,3,4). For analysis of the rate of pilus production per cell, single isolated cells were selected to ensure that no pilus is obstructed by a nearby cell. Otherwise there was no special selection. To increase statistical significance of the analyzed data, we used bootstrapping. In brief, for a set of analyzed data (e.g. pilus length) with *N* datapoints, we randomly picked *N* datapoints with replacement and analyzed this randomly picked dataset, e.g., by calculating the histogram. This process was repeated 10,000 times and statistical quantities of this bootstrapped data were calculated, e.g., the mean of all histograms and its 95 % confidence interval. Unless stated otherwise, P-values of two independent measurements were calculated using a two-tailed Wilcoxon-Mann-Whitney rank test in respect of the exponential shape of most distributions.

### Model description and estimation of rate constants

We propose a minimal 3-state model where the basal body of the motor can switch between three states: unbound, bound to the extension motor, and bound to the retraction motor. Both motors bind to the empty base in a mutually exclusive manner, with a binding rate of *k_ext,on_* and *k_ret,on_*, respectively. Binding of the extension motor leads to pilus extension and binding of the retraction motor leads to pilus retraction when a pilus is present. Both motors associated with the base unbind with rates *k_ext,off_* and *k_ret,off_*, respectively. A key assumption of our model is that all rates are independent of the piliation state. Thus, the retraction motor could bind to the base in a non-piliated state, in which case, it blocks the extension motor from binding. Binding or unbinding of a protein to a substrate are described by a Poisson process. In brief, the probability *Q*(*t*) that the binding/unbinding process of rate *k* does not occur until the time *t* has passed is described by the differential equation *Q*(*t*+*dt*) = (1 – *k dt*) *Q*(*t*) with the initial condition *Q*(*t* = 0) = 1, which yields the exponential decay *Q*(*t*) = exp(-*kt*). Therefore, the probability density to observe a binding/unbinding event at time *t*, is given by *P*(*t*) = *kQ*(*t*) = *k* exp(-*kt*). If the experimental time resolution *t*\* is finite, events shorter than *t*\* are missed which results in the shifted distribution 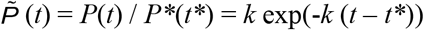 for *t* > *t*\*, where 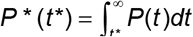. Due to our sampling rate of 20 Hz which is fast compared to the dynamics of the pilus, this shift is negligible except for the detection of multiple extension or retraction events (Supplementary Figure 2).

#### Binding rates of the motors

The binding rates of both proteins defines the dwell time between extension stalls and the subsequent extension or retraction starts. To calculate *k_ext,on_* and *k_ret,on_*, we looked into all identifiable pauses (extension-extension, extension-retraction, retraction-retraction) and their dwell times. In our model, those pauses correspond to an unbound state of the base. The probability that an extension/retraction motor binds at time *t* after the pause begins is given by *P_ext/ret_*(*t*) = *k_ext/ret,on_* exp(-*k_on t_*), where *k_on_* = *k_ext,on_* + *k_ret,on_*. The frequency of observing extension-extension events is thus given by 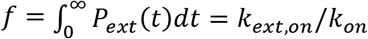. Due to limited resolution, we can only observe extension-extension events with an intervening dwell time *t* > *t*\* = 0.25 s. To include this limitation in our analysis, we estimate the ratio of observed extension-extension to extension-retraction events to be 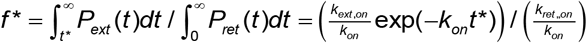. Experimentally, we obtain 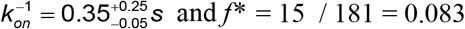, and consequently 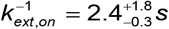 and 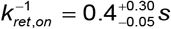.

#### Unbinding rate of the extension motor

In order to obtain *k_ext,off_*, we looked at the duration *T_ext_* of all pilus extension events. The model predicts that the probability of observing an extension event lasting for *t* = *T_ext_* follows an exponential distribution *P*(*t*) = *k_ext,off_* exp(-*k_ext,off t_*) (Fig. 2b). By fitting the exponential distribution to our experimental data, we obtained 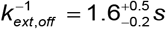. We further tested the hypothesis that the data are drawn from an exponential distribution using a one-sample Kolmogorov-Smirnov (K-S) test, which yields *P* = 0.87 indicating that the data are not significantly different from an exponential distribution, as expected.

#### Unbinding rate of the retraction motor

To obtain a meaningful estimation of *k_ret,off_*, we took into account the entire data set of both partial and full retraction events, and computed the maximum likelihood estimate of *k_ret,off_* that maximizes the likelihood of observing all the events. The probability that the retraction motor stays attached to the base after a pilus becomes fully retracted in time *t_full_* is *P_full_*(*t_full_*) = exp(-*k_ret,off_ t_full_*). The probability to observe a partial retraction where the retraction motor unbinds within a time window *t_part_* – *Δt* to *t_part_* is given by *P_part_*(*t_part_*) = exp(-*k_ret,off_*(*t_part_* – *Δt*)) – exp(-*k_ret,off_ t_part_*), where *Δt* 0.1 s is the finite time resolution of the experiment. The likelihood *L* of observing the partial and full retraction events 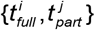 presented here as a function of *k_ret,off_* is then given by
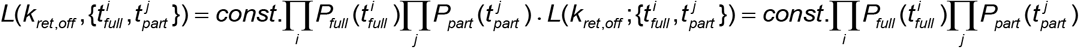. As shown in Supplementary Figure 3, we varied *k_ret,off_* and found the maximum of the likelihood function at 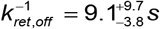. Errors represent the e^−1^ drop of the maximum likelihood.

### Monte Carlo simulation (MCS) of the dynamic binding of the motor proteins

To simulate the dynamic binding and unbinding of both proteins, we used the Gillespie algorithm^51^. In brief, the algorithm tracks the three variables: time *t*, pilus state *S*, and pilus length *L*. The states with the extension motor bound, the retraction motor bound, and empty base are denoted, respectively, by *S* = *E*, *S* = *R*, *S* = *ϕ*. The algorithm updates the variables as follows:

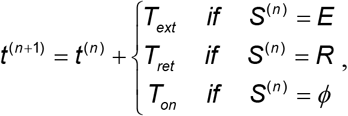

where *T_ext_*, *T_ret_*, and *T_on_* are randomly chosen from the exponential distributions defined by *k_ext,off_*, *k_ret,off_*, *k_on_*;

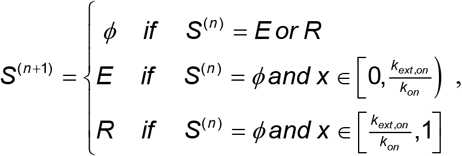

where *x* is a random number drawn from a uniform distribution on the interval [0, 1], and

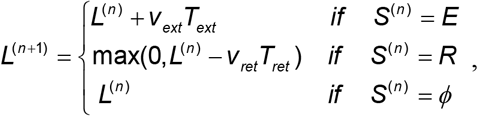

where for each step, the extension/retraction speed *v_ext_* = 485 ± 170 nm/s and *v_ret_* = 750 ± 314 nm/s are randomly chosen from a normal distribution with mean ± standard deviation as described in the main text. We then simulated 10^4^ 100-minute long trajectories of the stochastic model with the rate parameters taking the mean, lower bound, or upper bound values of the experimental measurements. From these simulated data, we first calculated if retraction resulted in a partial of full retraction event. Then, we calculate the distributions and their 95 % confidence intervals for all quantities and added them to the corresponding figures (Fig. 1e for pilus length, Fig. 2b for extension time, and Figs. 2d and 3b for retraction time). For each simulated trajectory, we then randomly cut out a one-minute long trajectory and counted the number of pili made in that one minute time window (see Fig. 3e).

To test whether the experimentally obtained distribution of pilus production rate is consistent with the simulated distributions of the three model, we performed a Kolmogorov-Smirnov test. Corresponding to the experimental data size, we computed the production rate for 10000 sets of 111 pilus machines in simulations and calculated the distribution of the distance *D* between the cumulative distribution of each individual simulation to the average of all simulations. We then measured the distance *D*_exp_ of the experimental distribution of pilus production rates to the same average of the simulation. This allowed us to obtain a measure of the statistical significance of our experimental data. Specifically, we defined the P-value as the probability of *D* > *D*_exp_. Here, the null hypothesis is that the experimental data is generated by the particular model. Consequently and opposite to a standard T-test, a large P-value (P > 0.05) means the null hypothesis cannot be rejected and hence is a good fit between the data and the simulation while a small P-value (P < 0.01) indicates that the simulation is unlikely to reproduce the experimental data. To include the uncertainty of the experimental measurements of binding and unbinding rates, we calculated P-values for each model using the mean of the estimated rates, the lower bounds and the upper bounds. For Model 1, we obtained P < 1e-6, P = 0.02, P = 0.40 for the lower bound, the mean, and the upper bound of the rates respectively. For Model 2 and 3 we obtained P < 1e-6 for all combinations of rates. For the ease of reading, we converted these P-values to regular P-values in the main manuscript, i.e., P < 1e-3 and P > 0.05 of the Kolmogorov-Smirnov test are converted to P > 0.05 and P < 0.001, respectively, in the main manuscript text.

## Supporting information

Sypplementary Movie 1

Sypplementary Movie 2

Sypplementary Movie 3

Sypplementary Movie 4

Sypplementary Movie 5

Sypplementary Movie 6

Sypplementary Movie 7

Sypplementary Movie 8

Sypplementary Movie 9

Sypplementary Movie 10

Sypplementary Movie 11

## Acknowledgements

We gratefully acknowledge invaluable help, discussions, and comments on the manuscript from Benjamin Bratton, Geoff Vrla, Joseph Sanfilippo, Nick Martin, Courtney Ellison, Benedikt Sabass, and support by our lab manager Joseph Sheehan. Additionally, we would like to thank Nicolas Biais and Jingbo Kan for initial advice on pilus labeling, Sagar Setru for implementing the ring-HiLo imaging at the optical trap and the patient help by Gary Laevski from the Princeton Molecular Biology Confocal Microscopy Core Facility and Nikon Center of Excellence

This work was supported by grants DP1AI124669 (to Z.G.), R21AI121828 (to J.W.S), and R01GM082938 (to N.S.W.) from the National Institute of Health, grant PHY-1806501 (to J.W.S.) from the National Science Foundation, fellowship KO5239/1-1 (to M.D.K.) from the German Research Foundation DFG, and the Princeton Center for the Physics of Biological Function sponsored by the National Science Foundation grant PHY-1734030.

## Author Contributions

All authors designed the research. M.D.K. performed all experiments. M.D.K. and C.F. analyzed the data and performed simulations. C.F. and N.S.W. developed the mathematical model. M.D.K., J.W.S. and Z.G. wrote the paper.

## Competing Interests statement

The authors declare no competing financial interest.

