## Supplementary Information for "Competitive binding of independent extension and retraction motors explains the quantitative dynamics of type IV pili"

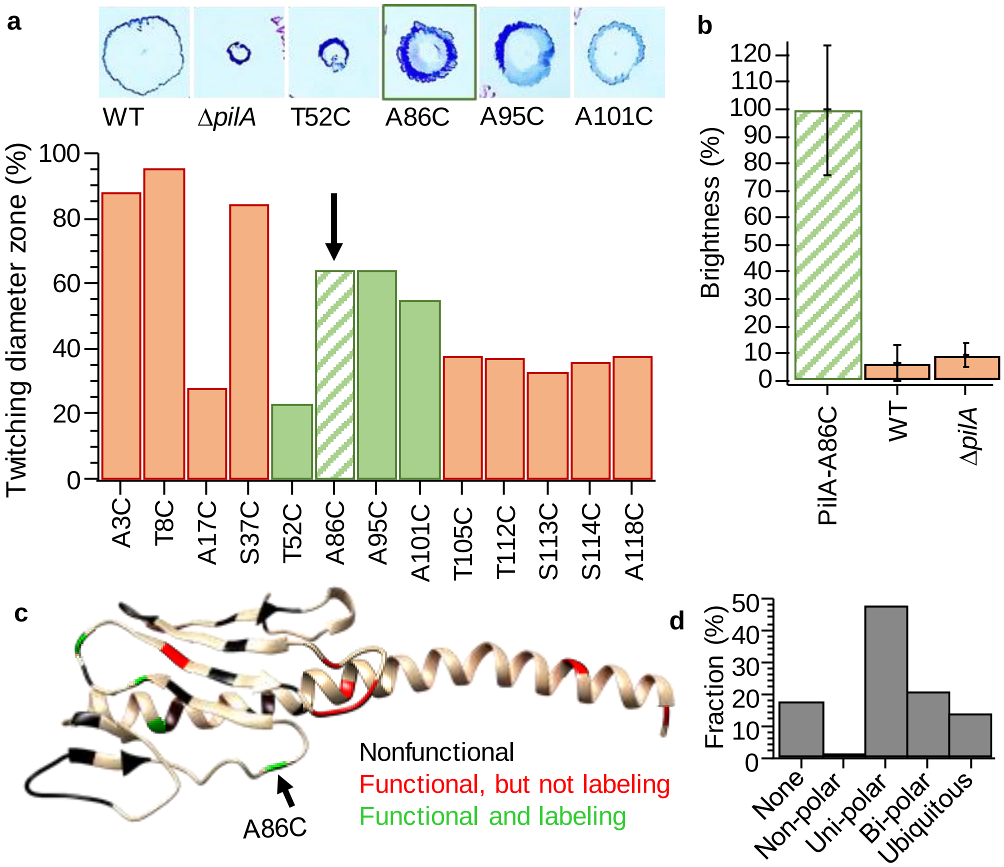


**Supplementary Figure 1 | PilA point mutations and their degrees of functionality. a,** The degree of functionality was determined by a standard twitching assay. In brief, single colonies grown overnight on agar plates were inoculated with the tip of a pipet between the plastic of a petri dish and 1.0% LB agar. After three days of incubation at 30 ºC, agar was carefully removed from the dish and colonies were stained with 0.1 % Crustal Violet for 10 minutes (top inset). The diameter for each mutant $D_m$ was measured and the relative diameter of the twitching zone $D_f$ (bottom) was estimated relative to the diameter of WT $D_{WT}$ and Δ*pilA* $D_{pilA}$ according to $D_f = (D_m - D_{pilA}) / (D_{WT} - D_{pilA})$. Mutations that resulted in labeled pili (by Alexa488-mal) are marked in green (T52C, A86C, A95C, A101C) while mutations that did not result in labeled pili are shown in red (A3C, T8C, A17C, S37C, T105C, T112C, S113C, S114C, A118C). For reference, the following point mutations resulted in non-functional pili: A36C, S41C, A44C, A45C, T51C, A62C, T70C, T71C, A72C, S73C, T74C, A75C, T76C, T78C, A93C, S99C, T107C, T110C, T119C, T123C, T127C, A128C, C134A, C134KO, C134S, C134T, S136C, T137C, C147A, C147KO, C147S, C147T. **b,** Labeling of PilA-A86C with Alexa488-mal is highly specific as WT and Δ*pilA* strains have 10 – 20 fold decreased brightness of the cell body compared to the PilA-A86C strain. **c,** Protein structure of PAO1 PilA obtained from the Phyre2 website and visualized with the Chimera software. **d**, Localization of pili in individual cells obtained by analyzing 30-second long movies.

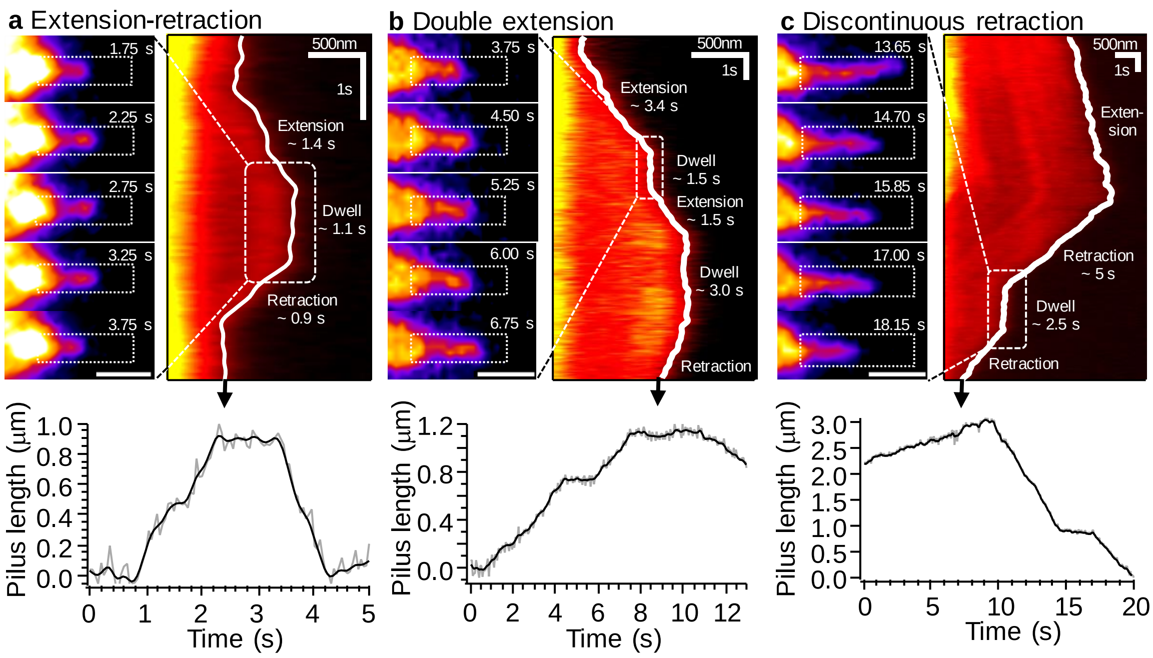


**Supplementary Figure 2 | Estimation of pilus length as a function of time. a,** Typical extension-retraction event. **b**, Double extension event. **c**, Discontinuous retraction event. **a,b,c** Individual frames of a movie (left panels) were analyzed by drawing a wide line scan (dotted white box) along the pilus. The fluorescence intensity of this line scan was plotted as a function of time in a kymograph (right panels). The pilus tip was then tracked using thresholding of the kymograph. The resulting trajectories (gray lines) of pilus length (lower panels) were smoothed slightly to reduce noise (black lines).

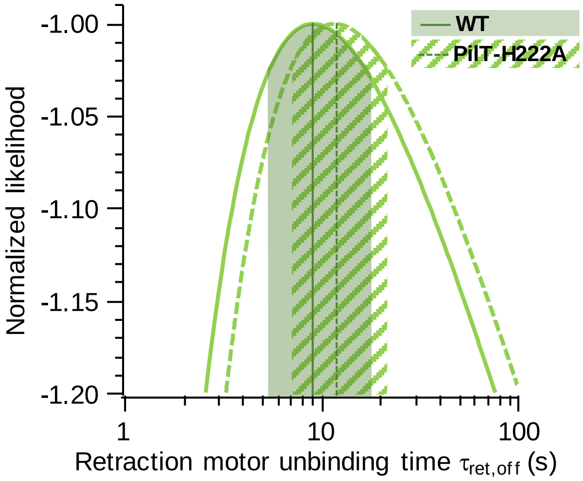


**Supplementary Figure 3 | Maximum likelihood approximation of PilT off rate and comparison between WT and slowly retracting mutant PilT-H222A.** Normalized likelihood of the observed partial and full retraction events, plotted against the characteristic time parameter $\tau_{ret,off}$ of unbinding of the retraction motor (see Methods for details). Shaded regions represent the 95% confidence interval.

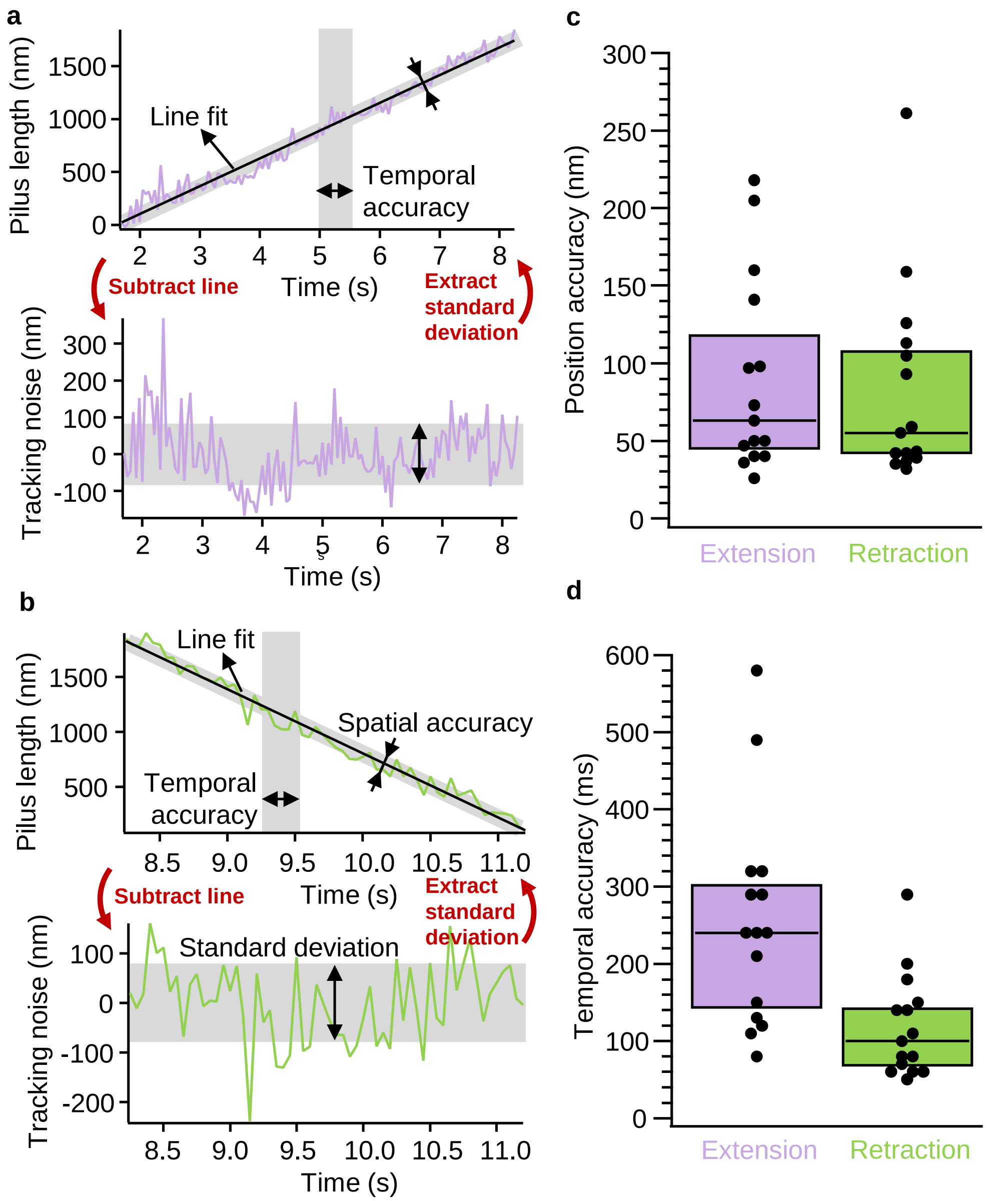


**Supplementary Figure 4 | Precision of pilus tracking. a**, Individual pilus extension and **b**, individual pilus retraction events are fitted with a line to obtain the trajectory of extension/retraction. This line was then subtracted from the data to find the positional tracking noise component of the data. As a measure for the average positional noise in the dataset, the standard deviation was estimated. The temporal uncertainty was estimated from the positional standard deviation by dividing by the slope of the pilus extension/retraction line fit. **c**, Distribution of positional accuracy. **d**, Distribution of temporal accuracy. This analysis assumes that extension and retraction velocities are constant and that all noise is imaging noise.

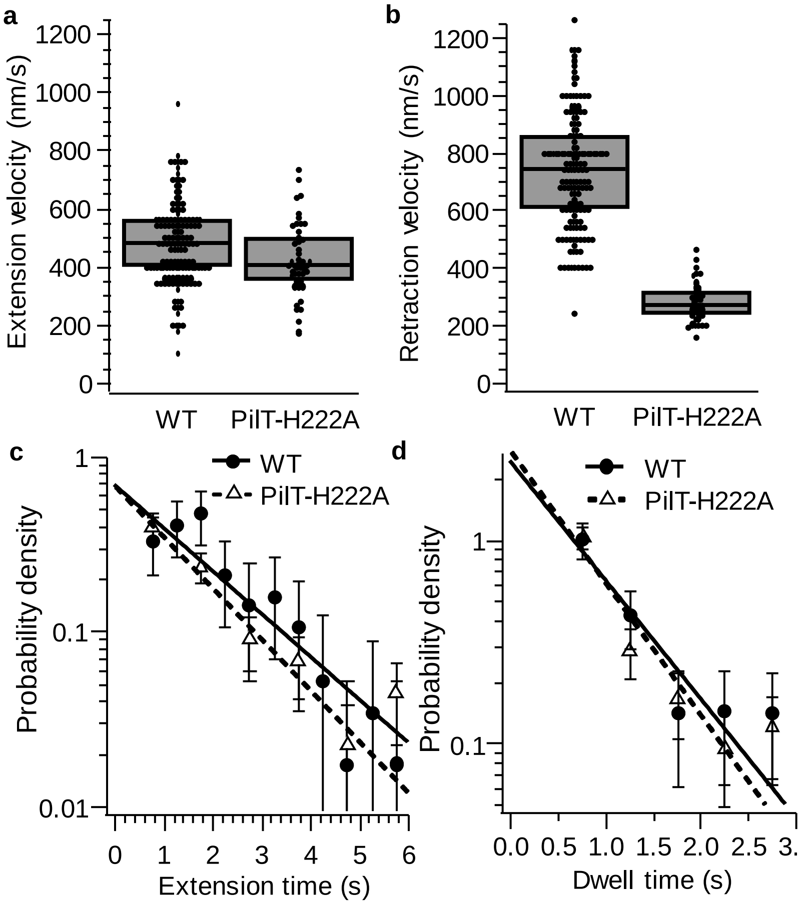


**Supplementary Figure 5 | Comparison of model parameters for WT and the slowly retracting mutant PilT$^{H222A}$. a**, Box plot of extension velocities with 25%/75% quantiles and median. **b**, Box plot of retraction velocities with 25%/75% quantiles and median. **c**, Distribution of extension times with exponential fits. **d**, Distribution of dwell times with exponential fits. (See Supplementary Figure 3 for comparison of the inferred unbinding rates of the extension motor in both backgrounds).

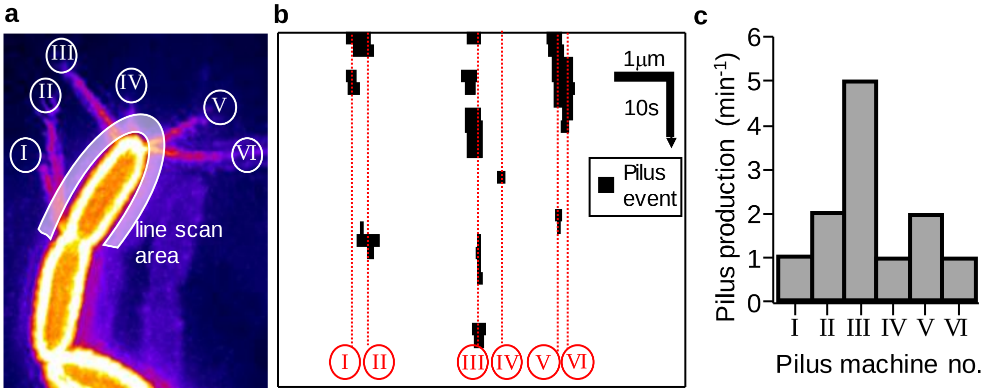


**Supplementary Figure 6 | Experimental estimation of pilus production rate of individual pilus machines (labeled I – VI). a**, Maximum projection of a super resolution (SIM) movie stack. Pili made by different machines that are close by are extended under different angles (e.g., I and II). A thick line scan (white transparent) close to the cell body is used to analyze intensity changes over time indicating pilus extensions. **b**, Kymograph of the binarized line scan intensity from a. White indicates background intensity, white indicates the intensity corresponding to a pilus crossing the line scan. Red lines indicate the positions of individual pilus machines as identified in a. **c**, Histogram of the number of individual pilus extension events counted in b.

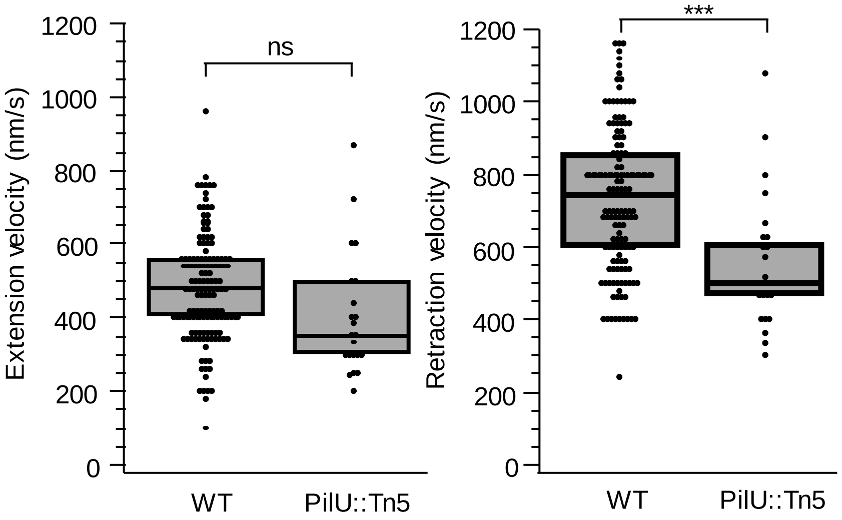


**Supplementary Figure 7 | Pilus extension velocity in a PilU mutant is indistinguishable from WT but PilU mutants retract 1.5 times slower than WT.** (ns: not significant; ***: P < 0.001)

| Strain | Description | Reference |
|---|---|---|
| **E. Coli** | | |
| S17 | Wild-type, used for cloning and conugation | |
| **P. aeruginosa** | | |
| PAO1 | Wild-type from Manoil transposon mutant library | (1) |
| ZG 1586 | *pilA-A86C* | This study. |
| ZG 1593 | Δ*fliC* | This study. |
| ZG 1587 | Δ*fliC pilA-A86C* | This study. |
| PW1728 | *pilT* ::[*Tn5* ] | (1) |
| ZG 1589 | *pilT* ::[*Tn5* ] *pilA-A86C* | This study. |
| ZG 1590 | *pilT* ::[*Tn5* ] *glmS* ::[*Ptet* ::*pilT-H222A-Flag* ] *pilA-A86C* | This study. |
| PW1731 | *piU* ::[*Tn5* ] | (1) |
| ZG 1592 | *pilU* ::[*Tn5* ] *pilA-A86C* | This study. |

**Supplementary Table 1 | Strains used in this study.**

| Plasmid | Descrption | Reference |
|---|---|---|
| pUC18-mini-Tn7T-LAC | Integrates downstream of PAO1 *glmS* | (2) |
| pTNS2 | Helper plasmid for pUC18-mini-Tn7T-LAC | (3) |
| pEXG2 | Vector for generating deletion mutants and point mutants | (3) |
| pEXG2-pilA-A86C | Introduce A86C point mutaiton in PAO1 *pilA* | This study. |
| pEXG2-ΔfliC | Deletes *fliC* in PAO1 | This study. |
| pTN7-Ptet::mKate2 | Targets *mKate2* with tetracyclin inducible promoter to *glmS* | This study. |
| pTN7-Ptet::pilT-H222A | Targets *pilT-H222A* with tetracyclin inducible promoter to *glmS* | This study. |

**Supplementary Table 2 | Plasmids used in this study.**

| Primer | Sequence | Reference |
|---|---|---|
| pEXG2_Ver1 | GTTGCATGGGCATAAAGTTGCC | This study. |
| pEXG2_Ver2 | CGGGTCCTCAACGACAGG | This study. |
| pilA-A86C P1 | GATACAAAGCTTGCTGCCAAATCGAGGAAATCC | This study. |
| pilA-A86C P2 | TCGGCGTCGAGCCGGATTGTAACAAGTTGGGTGTAATTGCTGTAG | This study. |
| pilA-A86C P3 | CTACAGCAATTACACCCAACTTGTTACAATCCGGCTCGACGCCGA | This study. |
| pilA-A86C P4 | GATACAAAGCTTCCAAGGATGTCAGGGCCG | This study. |
| DfliC P1 | GATACAAAGCTTGCGACTGGATGCTCGAAGG | This study. |
| DfliC P2 | GGCTTAGCGCAGCAGGCTGTTGACTGTAAGGGCCATGG | This study. |
| DfliC P3 | CCATGGCCCTTACAGTCAACAGCCTGCTGCGCTAAGCC | This study. |
| DfliC P4 | GATACAAAGCTTCACTGTGTACGTCGTCGAGC | This study. |
| pMK47_F1.For | AAGCTAATTCGATCATGCATTAAGACCCACTTTCACATTTAAGTTGTTTTTCTAATCC | This study. |
| pMK47_F1.Rev | CTCACCATTTTCATTTTAGCTTCCTTAGCTCCTGAATTCCT | This study. |
| pMK47_F2.For | AAGCTAAAATGAAAATGGTGAGCGAGCTGATTAAG | This study. |
| pMK47_F2.Rev | GGGGATCCACTAGTGAGTCAAGTCTTCGCGATGATTCTGTG | This study. |
| pMK47_BB.For | CTCACTAGTGGATCCCCCGG | This study. |
| pMK47_BB.Rev | TTTCATTTTAGCTTCCTTAGCTCCTGAATTCC | This study. |
| pMK73_F1.For | AGCTAAGGAAGCTAAAATGAAAATGGATATTACCGAGCTGCTCG | This study. |
| pMK73_F1.Rev | GGGGATCCACTAGTGAGTCACTTGTCGTCATCGTCTTTGTAGTCGA | This study. |
| pMK73_Flag | CTTGTCGTCATCGTCTTTGTAGTCGAAGTTTTCCGGGATCTTCGC | This study. |
| PilT_H222A P2 | CAGGGTGCCGAATACCAGGGCGCCGGTCTCCGCCGC | This study. |
| PilT_H222A P3 | GCGGCGGAGACCGGCGCCCTGGTATTCGGCACCCTG | This study. |

**Supplementary Table 3 | Primers used in this study.**

| Figure | Panel | Biological replicate name | Number of cells | Number of pili/events | Framerate (Hz) | Technique |
|---|---|---|---|---|---|---|
| 1 | c | WT 2Hz D1 Field 1 | 30 | 30 | 2 | HiLo |
| | | WT 2Hz D1 Field 2 | 20 | 19 | 2 | HiLo |
| | | WT 2Hz D1 Field 3 | 16 | 13 | 2 | HiLo |
| | | WT 2Hz D1 Field 4 | 20 | 13 | 2 | HiLo |
| | | WT 2Hz D1 Field 5 | 19 | 14 | 2 | HiLo |
| | | WT 2Hz D1 Field 6 | 22 | 17 | 2 | HiLo |
| | | WT 2Hz D2 Field 1 | 35 | 29 | 2 | HiLo |
| | | WT 2Hz D2 Field 2 | 42 | 35 | 2 | HiLo |
| | | WT 2Hz D2 Field 3 | 26 | 21 | 2 | HiLo |
| | | WT 2Hz D2 Field 4 | 27 | 24 | 2 | HiLo |
| | | WT 2Hz D2 Field 5 | 29 | 19 | 2 | HiLo |
| | | WT 2Hz D3 Field 1 | 49 | 40 | 2 | HiLo |
| | | WT 2Hz D3 Field 2 | 44 | 41 | 2 | HiLo |
| | | WT 2Hz D3 Field 3 | 28 | 22 | 2 | HiLo |
| | | WT 2Hz D3 Field 4 | 70 | 65 | 2 | HiLo |
| | | WT 2Hz D3 Field 5 | 65 | 58 | 2 | HiLo |
| | | Total | 542 | 460 | | |
| | d, e | WT 2Hz D1 | 16 | 89 | 2 | HiLo |
| | | WT 2Hz D4 | 78 | 267 | 2 | HiLo |
| | | WT 2Hz D5 | 28 | 150 | 2 | HiLo |
| | | WT 2Hz D6 | 74 | 526 | 2 | HiLo |
| | | Total | 102 | 676 | | |
| | f | WT 2Hz D1 | 3 | 4 | 2 | HiLo |
| | | WT 2Hz D4 | 16 | 25 | 2 | HiLo |
| | | WT 2Hz D5 | 15 | 40 | 2 | HiLo |
| | | WT 2Hz D6 | 32 | 74 | 2 | HiLo |
| | | Total | 47 | 114 | | |
| 2 | b | WT 20Hz D1 | 100 | 78 | 20 | HiLo |
| | | WT 20Hz D1 | 48 | 40 | 20 | HiLo |
| | | Total | 148 | 118 | | |
| | c | WT 20Hz D1 | 100 | 85 | 20 | HiLo |
| | | WT 20Hz D1 | 48 | 42 | 20 | HiLo |
| | | Total | 148 | 127 | | |
| | d | WT 20Hz D1 | 100 | 100 | 20 | HiLo |
| | | WT 20Hz D1 | 96 | 96 | 20 | HiLo |
| | | Total | 196 | 196 | | |
| 3 | a, b | WT 20Hz D1 | 100 | 100 | 20 | HiLo |
| | | WT 20Hz D1 | 96 | 96 | 20 | HiLo |
| | | Total | 196 | 196 | | |
| | | PilT-H222A D1 | 29 | 34 | 20 | HiLo |
| | | PilT-H222A D2 | 16 | 19 | 20 | HiLo |
| | | Total | 45 | 53 | | |
| | g | WT SIM D1 | 18 | 54 | 1 | SIM |
| | | WT SIM D1 | 16 | 57 | 1 | SIM |
| | | Total | 34 | 111 | | |
| 4 | d | WT 2Hz D4 | 10 | 42 | 2 | HiLo |
| | | WT 2Hz D5 | 14 | 65 | 2 | HiLo |
| | | WT 2Hz D6 | 7 | 43 | 2 | HiLo |
| | | Total | 21 | 108 | | |
| | | Trap D1 | 6 | 6 | 4 | R-HiLo |
| | | Trap D2 | 22 | 64 | 4 | R-HiLo |
| | | Total | 28 | 70 | | |
| | e | WT 20Hz D1 | 100 | 85 | 20 | HiLo |
| | | WT 20Hz D1 | 48 | 42 | 20 | HiLo |
| | | Total | 148 | 127 | | |
| | | Trap D1 | 6 | 4 | 4 | R-HiLo |
| | | Trap D2 | 22 | 22 | 4 | R-HiLo |
| | | Total | 28 | 26 | | |
| | f | WT 2Hz D4 | 78 | 267 | 2 | HiLo |
| | | WT 2Hz D5 | 28 | 150 | 2 | HiLo |
| | | WT 2Hz D6 | 74 | 526 | 2 | HiLo |
| | | Total | 208 | 969 | | |
| | | Trap D1 | 6 | 7 | 4 | R-HiLo |
| | | Trap D2 | 22 | 81 | 4 | R-HiLo |
| | | Total | 28 | 88 | | |

**Supplementary Table 4 | Table summarizing samples sizes for each figure.**

**Supplementary Movie 1**

PAO1 pilA-A86C mutant twitching on a 0.5% agarose pad.

**Supplementary Movie 2**

A single WT cell making several short and one very long pilus. Compare to Figure 1a in main text.

**Supplementary Movie 3**

Example for a dwell event between extension and retraction. Compare to Figure 1g and Supplementary Figure 2a.

**Supplementary Movie 4**

Example for a pilus double extension event with a dwell in between. Compare to Figure 1h and Supplementary Figure 2b.

**Supplementary Movie 5**

Example of a pilus discontinuous retraction. Compare to Figure 1i and Supplementary Figure 2c.

**Supplementary Movie 6**

Single cell on 0.5% agarose pad twitches forward due to pilus retraction.

**Supplementary Movie 7**

Single cell on 0.5% agarose pad twitches forward due to pilus retraction.

**Supplementary Movie 8**

A single WT cell held in liquid 5 μm above the cover slip by an optical trap.

**Supplementary Movie 9**

A single WT cell swimming into the optical trap, making one short pilus on the right-hand side pole, and then swimming out of the trap.

**Supplementary Movie 10**

A single Δ*fliC* cell that is incapable of flagella mediated swimming motility stays in the optical trap for prolonged time, making several pili on both poles. Compare to Figure 2b,c.

**Supplementary Movie 11**

A single WT cells producing several pili by several individual pilus machines taken by Structure Illumination Microscopy. Compare to Figure 5d and Supplementary Figure 6.
